# Two distinct proprioceptive representations of voluntary movements in primate spinal neurons

**DOI:** 10.1101/247213

**Authors:** Saeka Tomatsu, Geehee Kim, Joachim Confais, Tomohiko Takei, Kazuhiko Seki

## Abstract

When willingly setting our body in motion, we simultaneously know where and how our limbs are moving. While this indicates that proprioceptive information is readily represented in the neurons of the central nervous system, it is still unclear how. We recorded the activity of spinal neurons with direct projections from muscle spindle afferents in four monkeys, while they performed simple wrist movements. Against the assumption that these spinal neurons act as a simple relay of afferent input, we found the majority (56%) of neurons had firing patterns incongruent with a simple representation of spindle activity, and the minority had congruent patterns. Two groups of neurons showed distinct intrinsic characteristics (spike width, base firing rate and firing irregularity), and distinct control of their input-output gain. These results are the first demonstration that proprioceptive representation is achieved by the coordinated activity of distinct groups of neurons during volitional movement.

## Introduction

Without observing them, we know where our limbs are and how they are moving. How this proprioceptive information is encoded in central nervous system (CNS) neurons has been a central question in sensorimotor control research (Gandevia, 1996; Proske & Gandevia, 2009). Muscle spindles are now widely accepted as being the primary receptor organs mediating proprioception (Edin & Abbs, 1991). The primary and secondary endings of the spindle encode muscle length, as well as its rate of change, as the core component of proprioception, and their passive length-response characteristics have been well established in experiments using anesthetized animals and healthy human subjects (Cheney & Preston, 1976; Nathan, Smith, & Cook, 1986; Poppele & Kennedy, 1974).

However, during voluntary movement fusimotor control prevents spindle activity from being explained solely by the length of the parent muscles (Dimitriou & Edin, 2008, 2010; Ribot, Roll, & Vedel, 1986). During a simple joint movement, for example, a limb muscle is shortened by agonistic, and lengthen by antagonistic, movements. If the descending drive to the intra-fusal fibers dominates the spindle afferent activity encoding the passive length of extra-fusal muscle fibers (Edin & Vallbo, 1990), spindle activity during active shortening could became greater than during passive shortening, or may even passively lengthen. Indeed, spindle activity has been shown to primarily represent the speed and acceleration, rather than the length, of parent muscles during human finger movements (Dimitriou & Edin, 2008, 2010). Nevertheless, spindle afferent recording in cats (Loeb, Hoffer, & Pratt, 1985) and monkeys (Schieber & Thach, 1985), as well as microneurography studies in human subjects (al-Falahe, Nagaoka, & Vallbo, 1990a, 1990b), suggest that changes in muscle length could override the effect of alpha-gamma motor neuron linkage. Therefore, there is a general agreement that muscle length information is dominantly encoded by spindle afferents, both at rest and during voluntary muscle contraction.

In the CNS, these spindle afferents project to postsynaptic neurons either in the brainstem or spinal cord. Under anesthesia, the features encoded in spindle afferents, including the length of parent muscles, are also represented in postsynaptic neurons (Mackie, Morley, & Rowe, 1999; Mackie, Morley, Zhang, Murray, & Rowe, 1998; Pompeiano, Wand, & Sontag, 1975; Wand, Pompeiano, & Fayein, 1980). However, in awake, behaving conditions, the transmission of spindle afferents to relay neurons in the CNS may not always be exact. Postsynaptic neurons of spindle afferents in both the spinal cord and cuneate nucleus also receive inputs from the cortex and descending tracts (Bentivoglio & Rustioni, 1986; Cheema, Rustioni, & Whitsel, 1985; Leiras, Velo, Martin-Cora, & Canedo, 2010; Noble & Riddell, 1989; Weisberg & Rustioni, 1977) and from multiple classes of primary afferents (Bosco & Poppele, 2001; Jankowska, 1992; Jankowska, Rastad, & Zarzecki, 1979; Oscarsson, 1973). During voluntary movement, these inputs could activate single neurons in various ways. In that case, how can information about muscle length, provided by spindle afferents, be represented in CNS neurons by overcoming these convergent inputs during voluntary movement?

To address this issue, we trained monkeys to perform a simple wrist flexion-extension movement, recorded the activity of spinal neurons postsynaptic to group Ia spindle afferents from wrist extensor muscles, and compared their activity during agonistic and antagonistic movements.

## Results

We analyzed the activity of cervical spinal neurons showing a response to electrical stimulation applied to proprioceptive deep radial (DR) nerves (pure muscle nerve innervating wrist extensor muscles) in four *Macaca sp.* Monkeys performing a wrist flexion-extension task (Figure 1; See materials and methods for detail). For comparison, we also recorded the neurons responding to stimulation of the cutaneous superficial radial nerve (SR, pure cutaneous nerve innervating the dorsoradial aspect of hand). Stimulation was applied through chronically implanted nerve cuff electrodes, during a wrist flexion-extension task. A total of 292 spinal neurons (50 from monkey KO, 47 from monkey IS, 63 from monkey OK, and 132 from monkey KJ) showed evoked responses within 10 ms after the stimulation of either nerve type. Of these neurons, 150 (38 from KO, 25 from IS, 50 from OK, and/or 37 from KJ) were classified as postsynaptic neurons from the SR (*n* = 57) and DR (*n* = 93) and six neurons responded to both nerves. After excluding from further analysis two neurons that were classified as putative motoneurons, the total number of neurons analyzed was 142. This dataset overlaps with a previous paper (Confais, Kim, Tomatsu, Takei, & Seki, 2017) that focused on the differences between SR and DR neurons (i.e. responsiveness to nerve stimulation, central latency, base firing rate, and intra-spinal depth of each neuron). In brief, SR neurons were located more superficially than DR neurons in the dorsal horn of the cervical spinal cord (p < 0.05); no difference was found in their base firing rate (p > 0.05) and threshold currents for DR and SR neurons were 1.16 ± 0.20 and 1.73 ± 0.65 T (times the threshold, mean ± SD), respectively. The SR neuron threshold current (< 2 times the threshold to evoke the incoming volley) suggests that afferents from cutaneous mechanoreceptors were mainly recruited to activate these neurons (Macefield, Gandevia, & Burke, 1989), as opposed to nociceptors. The lower threshold of DR neurons suggested that group Ia afferents were preferentially recruited over group Ib afferents by nerve stimulation (Bradley & Eccles, 1953).

**Figure 1.**
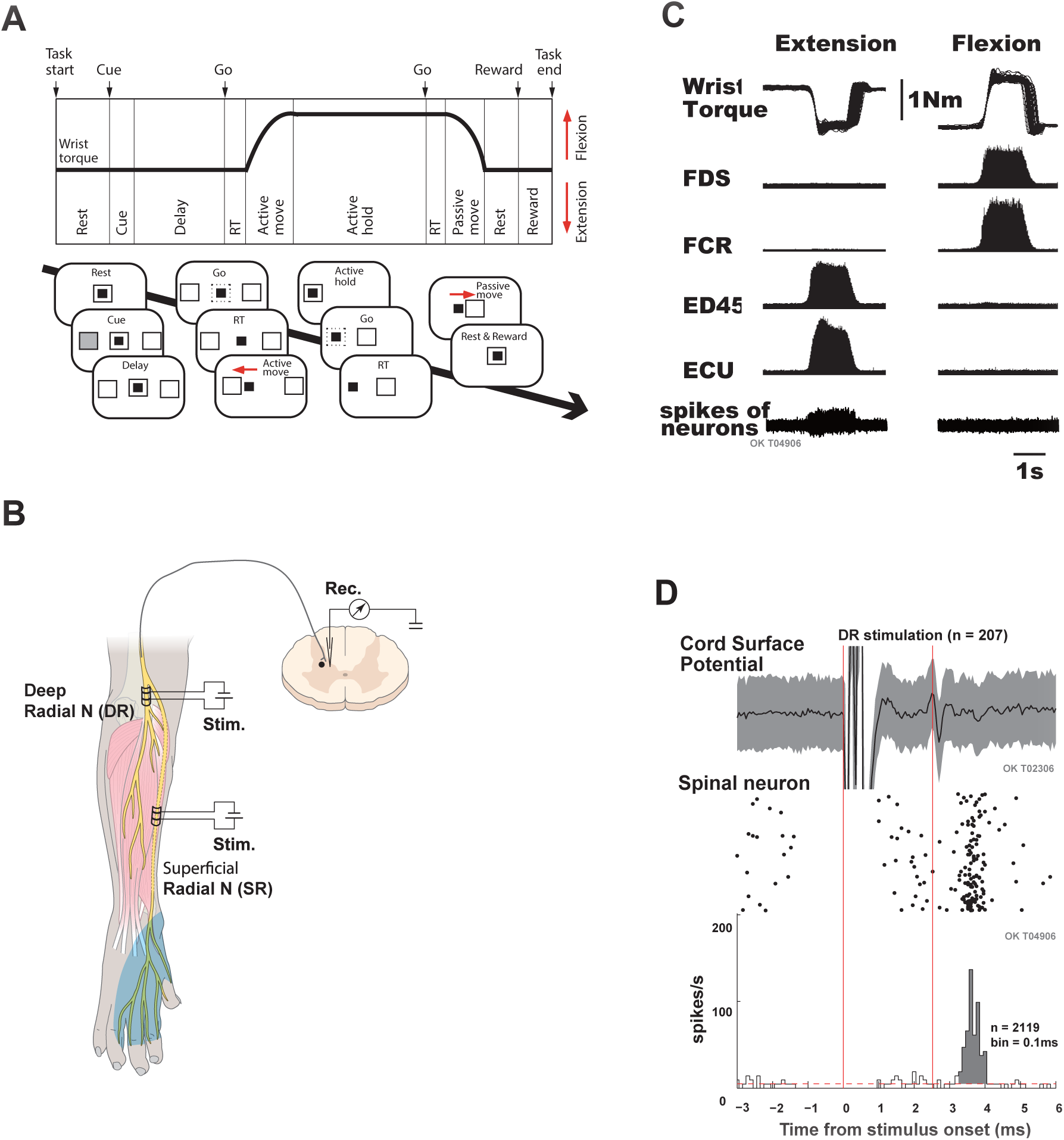
Experimental setup. (A) Task sequence. (B) Inputs from peripheral nerves. The radial nerve has two distal branches. The deep radial (DR) nerve provides sensory innervation to the wrist extensor muscles (red shade), and the superficial radial (SR) nerve provides it to the skin of the dorsal aspect of the thumb, index, and middle fingers and the radial half of the ring finger (blue shade). Nerve cuff electrodes were implanted into the DR, SR, and median nerve to classify the putative first-order interneurons in the spinal cord. Spinal neural activity was recorded using a metal microelectrode placed in monkeys performing wrist flexion and extension tasks. (C) Representative data of wrist torque (top), electromyographic activity (EMG) of wrist extensor (ECU, ED45) and flexor (FDS, FCR) muscles (middle), and single spinal interneuron (#OK4906) activity (bottom) while the monkey OK performed a wrist flexion and extension task. Wrist flexor and extensor muscles were alternately active during flexion and extension trials, respectively. This specific example of DR-neurons was exclusively activated during the extension trial. (D) Classification of putative first-order interneurons. *Top*; cord surface potential (average of 207 sweeps and area of standard deviation of individual sweeps) evoked by stimulation of the DR nerve (1.1–1.2 times the threshold). *Bottom;* Raster plot and peristimulus time histogram (PSTH) of the interneuron activity shown in *C*. Segmental latency was measured from the cord surface volley peak (red vertical line, 2.55 ms after stimulation) to the onset of the PSTH peak. The onset of the PSTH peak was defined as an increase of two standard deviations (red dashed line) from the mean line (−50 ms to −1 ms relative to stimulus onset). The segmental latency of this neuron was 0.65 ms, and the evoked response jitter from the onset and offset of the PSTH peak was 0.7 ms, indicating that this neuron was a DR neuron.

### Activity modulation of DR neurons during the wrist flexion–extension task

Examples of the activity of DR neurons during voluntary wrist flexions and extensions are shown in Figure 2A. Compared with flexion movements, the firing rate of this neuron was higher and more sustained during extension, where the parent muscle of DR afferents undergoes a shortening contraction (agonistic movement). Additionally, activity during extension only started earlier than the onset of active movement (Prut & Fetz, 1999). An SR neuron example is shown in Figure 2B. Activity was generally biased toward extension, but the difference between flexion and extension was not as dominant as it was for the DR neuron shown.

**Figure 2.**
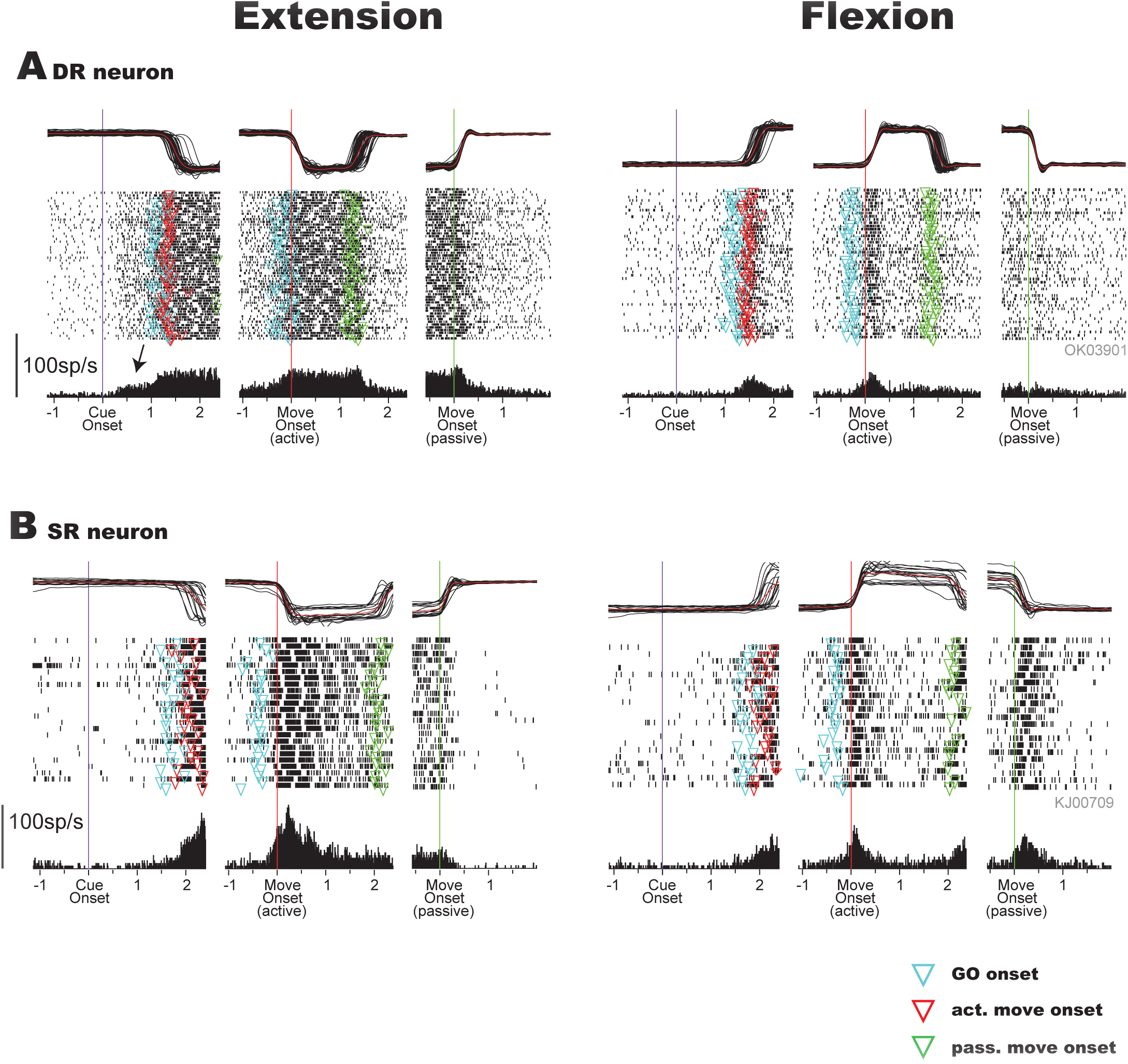
Examples of task-related activity of DR and SR neurons. Representative examples of DR neurons (top) and SR neurons (bottom) neurons before (left in each panel), during (center), and after (right) the active wrist movement were plotted separately for extension (left column) and flexion (right column) trials. In each panel, the time courses of mean (red) and trial-by-trial (black) smoothed extension–flexion (E–F) torque (top), raster plots for each trial (middle), and peri-event time histograms (PETH) were aligned at three different time points: onset of cue signal (purple lines), onset time of active movement (red lines and inverted red triangles), and onset time of passive movement (green lines and inverted green triangles). Inverted light blue triangles indicate onset of go signal. DR‐ and SR-neurons were recorded from monkeys OK and KJ respectively. Please note that the activity of neuron in A started after the onset of the cue signal but before the EMG onset (see a black arrow).

Next, we calculated the mean firing rate during each task epoch using data from all DR and SR neurons (Figure 3A and B). Many of the characteristics observed in the neurons in Figure 2 are reproduced in these plots. The median of the mean firing rate for five behavioral epochs clearly indicated that the DR and SR neurons increased their activity during active torque (active move and active hold) and passive movement for both flexion and extension movements. Their activity was significantly larger in extension trials (Wilcoxon signed rank test, active move, P = 0.001; active hold, P = 0.020; passive move, P = 0.043) (see also (Confais et al., 2017)). We further characterized this activity by separately counting the number of neurons showing significant modulation relative to rest (Figure 3C) in each behavioral epoch. More than 30% of neurons exhibited significant modulation in the delay period preceding movement onset (Fig. 3C: 31.9% of DR neurons, 42.1% of SR neurons). Because this instructed delay period was defined as the epoch preceding the onset of the earliest electromyographic activity (EMG) activity (i.e. 300 ms before movement onset; see Materials and methods), any activity recorded during this period is more likely to be driven by descending motor commands than by reafferent signals. During active torque, almost all neurons exhibited significant modulations in at least one movement direction (active move: 90.1% of DR neurons, 94.7% in SR neurons; active hold: 89.0% of DR neurons, 89.5% of SR neurons), indicating that these neurons were involved in the voluntary wrist movement. Moreover, during active torque, approximately 70% of neurons were modulated in both movement directions (71.4% of DR neurons, 80.7% of SR neurons), and approximately 80% of neurons exhibited directional differences in firing rates (82.4% of DR neurons, 73.7% of SR neurons). During the passive move period, 71.4% of DR neurons and 80.7% of SR neurons exhibited significant modulation. Overall, 95.8% of neurons showed significant modulation in at least one behavioral epoch.

**Figure 3.**
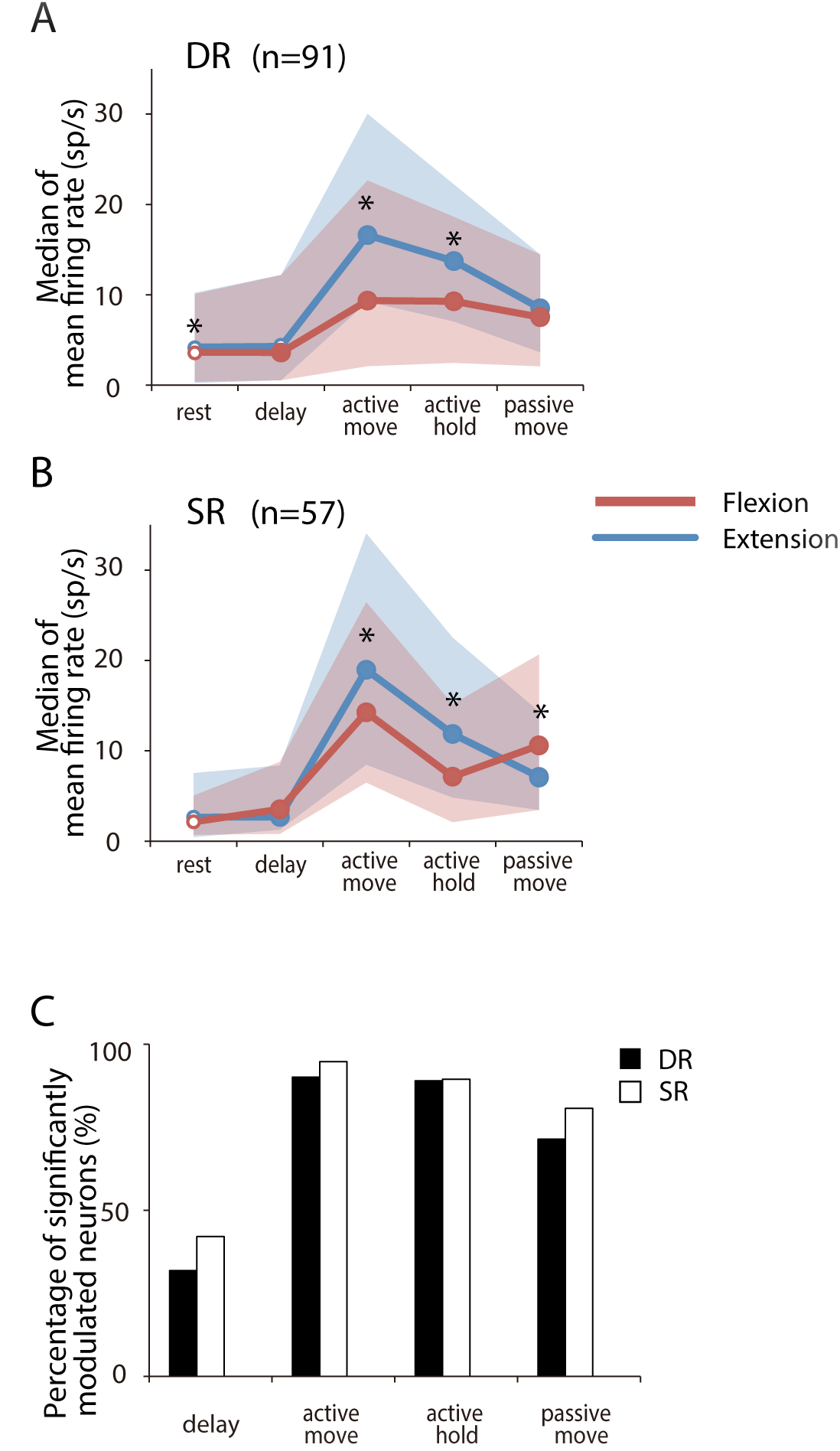
Population analysis of firing rate modulation of DR neurons. (A,B) Median of mean firing rate of five behavioral epochs for all neurons (DR, *n* = 91; SR, *n* = 57). Blue and red lines indicate extension and flexion trials, respectively. Blue and red shades indicate interquartile ranges. Small open circles indicate the mean firing rate in the epoch that did not exhibit significant modulation compared with the rest period (Wilcoxon signed rank test, *P*< 0.05). Asterisks indicate a significant difference between extension and flexion trials (Wilcoxon signed rank test, *P* < 0.05). (C) Percentage of significantly modulated neurons relative to the number of all neurons classified as DR or SR neurons. In this figure, we counted neurons that exhibited a significant modulation (suppression and facilitation from rest) in either one or both movement directions. There was no significant difference in the percentage of significantly modulated neurons according to the input nerve in each epoch (X-square test, P > 0.25).

### Two types of DR neurons with high and low fidelity to spindle inputs

As summarized in Fig. 3, the majority of DR neurons were predominantly active during active extension torque (Confais et al., 2017). This pattern of activity was paradoxical because during extension, extensor muscles (i.e. spindle-bearing muscles of DR afferents) are primarily shortening. Under the assumption that DR neurons are primarily activated by Ia afferents from their parent muscles, one could expect more activity in the direction that stretches their spindle-bearing muscles, that is, flexion (al-Falahe et al., 1990b; Dimitriou, 2014; Jones, Wessberg, & Vallbo, 2001).

We then addressed how these neurons represent the length of their parent muscles. Examination of Fig. 3 implies that as a population, DR neurons show non-negligible activity during active flexion, that is, when the parent muscle is stretched. This raises the possibility that the length of the parent muscle could be represented in the minority of DR neurons. We tested this hypothesis by selecting the DR neurons showing significantly more activity during active flexion than during active extension. To this end, we compared the mean firing rate during active torque (both active movement and hold) between extension and flexion trials in each DR neuron. We that found 23/91 DR neurons had flexion-biased activity. Examples of these neurons’ activities are shown in Figure 4. The neuron shown in A exhibited significant facilitation during flexion (paired t-test, P < 0.001) and significant suppression during extension (paired t-test, P < 0.001), both compared with during the rest period. This firing pattern is consistent with a representation of the length of the parent muscle (see illustration at the top) and was seen in 7/23 of these neurons. The neuron in B also had greater activity during active torque for both extension and flexion, but significantly larger in flexion than extension (paired t-test, P < 0.001). Again, this firing pattern is consistent with a representation of the length of the parent muscle and was seen in 11/23 of these neurons. Remarkably, this neuron also had dominant activity during the instructed delay period preceding the active shortening (i.e. before flexion movement). Similar facilitation during the instructed delay period preceding flexion was observed in 7/11 neurons in this group. Finally, in both neurons shown in A and B, the background firing rate (e.g. before cue onset) was clearly greater compared with typical DR neurons exhibiting greater activity during extension (C; see also Fig. 2A). As the neurons shown in A and B had a pattern of activity that one would expect from Ia afferents, we termed them “high fidelity neurons” (HFNs) in response to expected spindle afferent inputs. Unexpectedly only a minority (23/91) of DR neurons with Ia inputs represented the spindle-like activity. In contrast, the majority (51/91) of neurons exhibited more activity during extension, as exemplified in Figures 2A and 4C. Because this pattern of activity cannot simply be explained by the spindle activity of parent muscles, we termed these neurons “low fidelity neurons” (LFNs).

**Figure 4.**
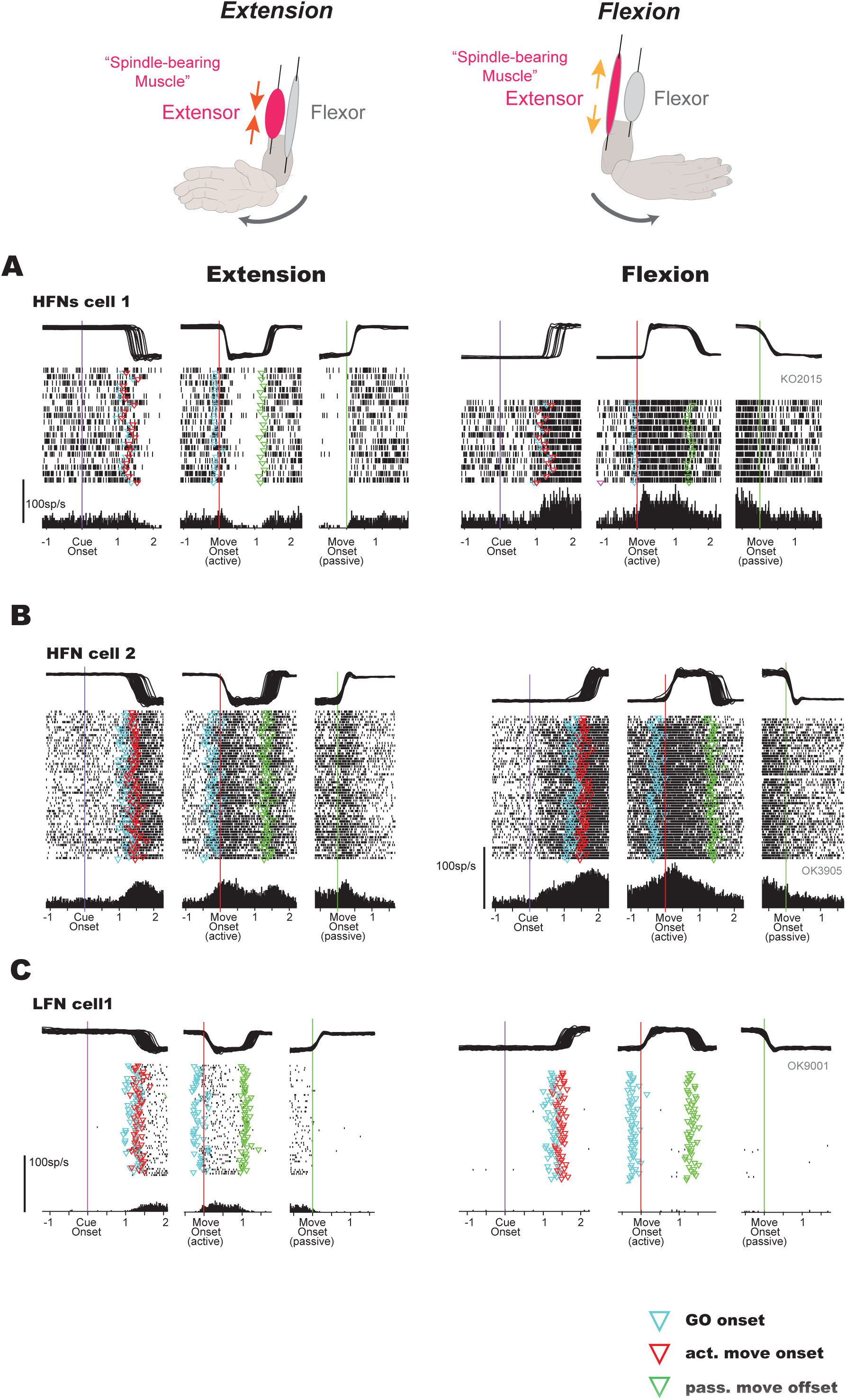
Example firing patterns of high fidelity neurons (HFN) and low fidelity neurons(LFN). Representative examples of DR neurons showing a firing pattern congruent with the putative spindle activity of their host muscle (HFN) (A, B), and of an LFN showing an incongruent pattern (C). See text for details. Format of each panel is the same as in Figure 2.

In addition, 11/91 neurons showed comparable activity between flexion and extension, and 6/91 DR neurons showed comparable activity between rest and active torque; the activity of these neurons could not be easily interpreted, and were excluded from further analyses.

The firing patterns of HFNs and LFNs are summarized in Figure 5. Greater mean firing rates during active flexion than extension in HFNs (Wilcoxon rank sum test, P < 0.001) are expected by definition. Surprisingly, however, we found that a greater HFN firing rate persisted during the majority of task epochs. Indeed, it overwhelmed that of LFN throughout the flexion task and during the period without movement. We thought this might indicate that HFNs and LFNs are represented from two distinct classes of spinal neurons with different intrinsic properties, so we examined this possibility.

**Figure 5.**
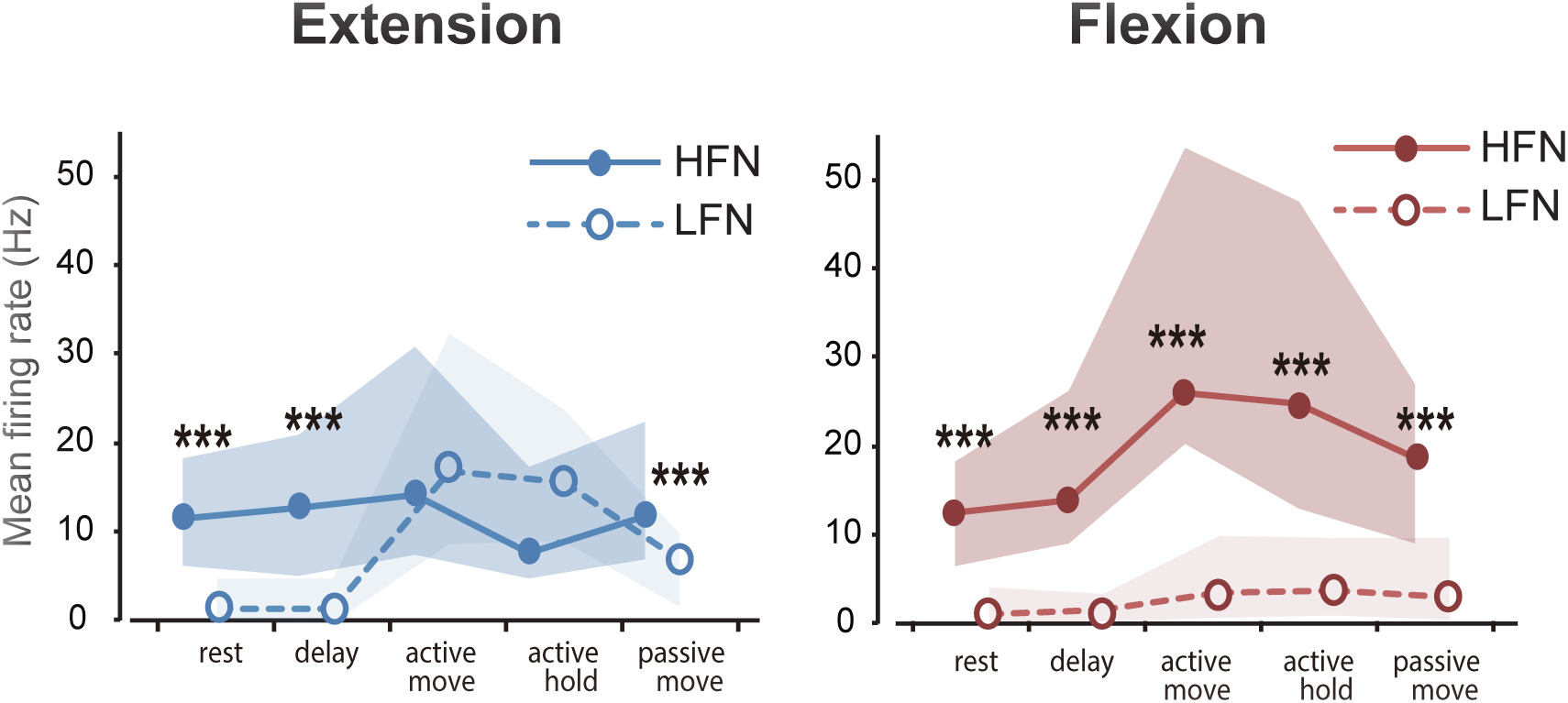
Population average firing pattern of HFNs and LFNs. Median of the mean firing rate in five behavioral epochs for two subgroups (HFNs (continuous line with filled circle; *n* = 23) and LFNs (dotted line with open circle; *n* = 51)) of DR neurons. In each DR neuron, the firing rate during active flexion and extension (both active movement and active hold period) was compared and categorized as HFN if larger during flexion, and as LFNs if larger during extension. Blue and red lines indicate extension and flexion trials, respectively. The shaded areas represent the interquartile range around median. Asterisks indicate a significant difference between LFN and HFN (Wilcoxon sum rank test, ***, *P* < 0.001). (C) Mean peak areas during rest periods at the beginning of the task are compared between LFN (open) and HFN (filled) groups (two-sample t‐ test; **, *P* < 0.01)

### HFNs and LFNs have different intrinsic firing properties

Because interneurons have shorter action potentials than projection neurons (Connors & Gutnick, 1990), and the extracellular waveform reflects the intracellular waveform (Henze et al., 2000), spike duration has been used to distinguish different class of neurons in the neocortex (Kaufman et al., 2010; Merchant, Naselaris, & Georgopoulos, 2008) (Miri et al., 2017; Mitchell, Sundberg, & Reynolds, 2007). Following these studies, we compared the spike duration of both groups, as a potential clue regarding their intrinsic properties in Figures 6A and D. The spike durations in both groups ranged between 0.12–0.53 ms, partially overlapping the range in the rodent (Bartho et al., 2004) and monkey (Kaufman et al., 2010; Merchant et al., 2008; Mitchell et al., 2007; Mountcastle, Talbot, Sakata, & Hyvarinen, 1969) cortices. Importantly, HFNs and LFNs had significantly different distributions. While spikes in most HFNs had short durations (< 0.25ms) (21/23, 91%), a sizable population of LFNs had longer durations (> 0.25ms) (18/51, 35%) alongside a majority of neurons with shorter durations (hatched area in open box, n = 33/51, 64%). On average, HFNs had significantly shorter durations than LFNs (two-sample t-test, P = 0.040c). This indicates that HFNs and LFNs are distinct classes of spinal neurons with different intrinsic properties; HFNs may be smaller interneurons with shorter action potentials, and LFNs may be larger projection neurons with longer action potentials.

**Figure 6:**
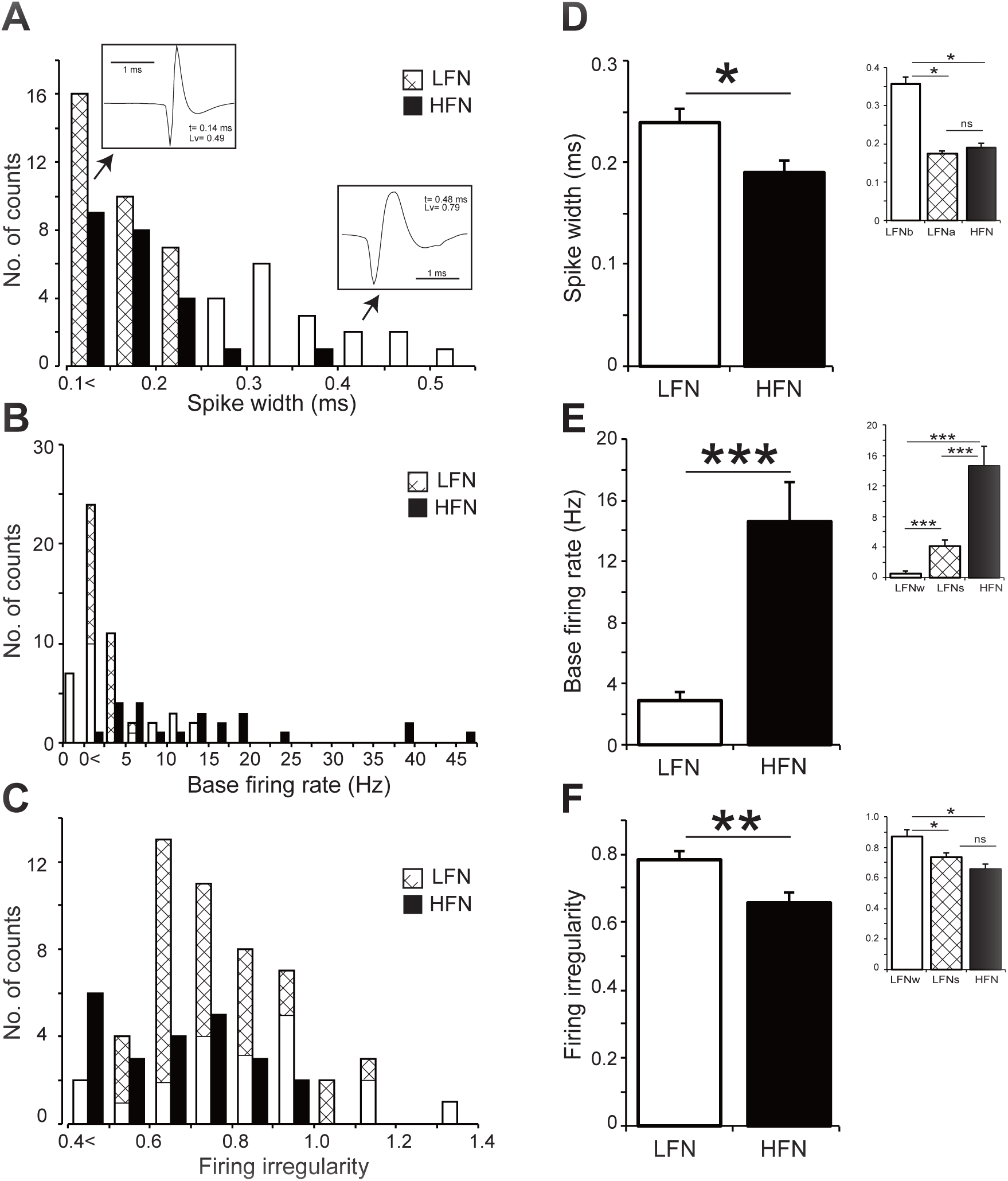
Comparison of spike width, base firing rate, and firing irregularity between HFNs and LFNs. A distribution (***A, B, C***) (A, B, C) and mean ± standard error (D, E, F) of spike-width (A, D), base firing rate (B, E), and firing irregularity (local variance [Lv] of inter-spike intervals (Shinomoto et al., 2003) (C, F) of LFNs (open) and HFNs (filled). Hatched area in open boxes correspond to LFNs with shorter action potentials (< 0.25 ms); and insets of panel D-F show the corresponding statistics. Asterisks indicate a significant difference between two groups (Wilcoxon rank sum test or two-sample t-test, \**P* < 0.05, ** *P* < 0.01, *** *P* < 0.001). Insets in panel A shows representative averaged waveforms of extracellular action potentials of HFNs (n = 8821, left) and LFN (n = 1603, right), respectively. The base firing rate and Lv of each neuron are also shown in each panel.

In the cerebral cortex, interneurons fire with higher spiking rates and more irregular spiking patterns (“fast spiking neurons”), while pyramidal neurons fire with smaller and more regular firing rates (“regular spiking neurons”) (Kaufman et al., 2010; Merchant et al., 2008; Miri et al., 2017; Mitchell et al., 2007; Mountcastle et al., 1969). Thus, we next compared the base firing rate and firing irregularity of HFNs and LFNs. Base firing rates were computed during the rest periods of both flexion and extension trials for each neuron. We found a dramatic difference between the two groups. While the base spontaneous firing rate in the majority of LFNs was < 5 Hz, the majority of HFNs were > 10 Hz (Fig. 6B). As a result, HFNs had significantly higher background firing rates than LFNs (Fig. 6E, (mean ± SD) 14.62 ± 12.54 vs 2.87 ± 3.85, Wilcoxon rank sum test, P < 0.001). Firing irregularity was evaluated by the local variation (Lv) of interspike intervals (ISIs) (Abbasi et al., 2017; Mochizuki et al., 2016; Shinomoto, Shima, & Tanji, 2003), and was found to be different between the two groups (Fig. 6C,F): LFNs tend to have larger Lvs than HFNs (0.78 ± 0.18 vs 0.65 ± 0.16, two-sample t-test, P = 0.005). In other words, HFNs fired more regularly (regular ISIs) than LFNs during the task.

It is important to point out that spike durations of a subgroup of LFNs tended to heavily overlap with those of HFNs (compared filled and hatched bars in A), and indeed was comparable with those of HFNs (insert in Fig. 6D). Both base firing rate (insert in Fig. 6E) and Lv (insert in Fig. 6F) also showed values closer to the HFNs. Therefore, this subgroup of LFNs might showing similar intrinsic properties to HFNs, while having a different activity pattern during flexion/extension. This suggests that intrinsic properties themselves are not the only distinction between HFNs and LFNs, and that other factors are needed to fully explain the differences in their firing patterns during wrist movement. Therefore, it does not seem likely that the differences in task-related activity between HFNs and LFNs can be solely ascribed to their respective intrinsic properties.

In contrast to data obtained from *in vitro* preparations or mathematical models (Connors & Gutnick, 1990; Mainen & Sejnowski, 1996), the firing characteristics of neurons recorded *in vivo* are not only determined by their intrinsic properties, but can also be influenced by their convergent synaptic inputs. For example, the higher background firing rate of HFNs could also be explained by a stronger synaptic drive. We explored this possibility by examining their input-output gains from primary afferents in the next section.

In addition, as with the other intrinsic properties of these neurons, we compared the putative recording depth of HFNs and LFNs using a method we previously reported (Confais et al., 2017; Takei & Seki, 2010). In short, for each recording track, we considered that the depth of the dorsal edge of the grey matter corresponded to that of the first encountered cell. This was then used as reference to gauge the depth of all cells recorded in the same track. The recording depth was (mean ± SD) 1414 ± 817 μm for the HFNs and 1514 ± 1118 μm for the LFNs, and no difference was found between these two groups (two-sample t-test, P = 0.634). In a previous study, we used the same technique to locate a spinal recording site at a depth of 1490 μm below the first cell (Takei & Seki, 2010). Subsequent histological analysis indicated that this site was located around the border of the dorsal horn and intermediate zone (Takei & Seki, 2010). Considering these data, we speculated that both group of neurons were most likely located in the anterior dorsal horn or the intermediate zone (laminae IV–VII) (Confais et al., 2017), with HFNs and LFNs being intermingled. The segmental latencies and threshold currents to recruit HFNs and LFNs were not significantly different between the two groups (segmental latency, 0.55 ± 0.42 ms (HFNs) and 0.60 ± 0.31 ms (LFNs); threshold current, 1.16 ± 0.22 T (HFNs) and 1.12 ± 0.09 T (LFNs)).

### Different input-output gains from spindle afferents to HFNs and LFNs

Previously we showed that input-output gains differ between rest and active torque for both DR neurons (Confais et al., 2017) and SR neurons (Confais et al., 2017; Seki & Fetz, 2003). We also found that different levels of input-output gain can be induced by different degrees of presynaptic inhibition to the primary afferent, in SR neurons at least (Seki, Perlmutter, & Fetz, 2003, 2009). Next, we examined the input-output gain of HFNs and LFNs from Ia afferents to investigate whether the modes of synaptic inputs, not just the intrinsic properties, are distinct between the two neuronal groups.

We compared the peak area of peristimulus time histograms (PSTHs) (Figure 1D) aligned to the stimulation of DR afferents as an index of the responsiveness of HFNs and LFNs during different behavioral epochs (Figure 7A, B). Responsiveness to DR stimulation was significantly larger in the HFNs than in the LFNs throughout the task, during both extension (A) and flexion (B) trials. This difference was particularly dominant during the rest period (C). As the base firing rates preceding nerve stimulation are subtracted from the PSTH peak value, such a result is not an artifact due to the difference in firing rate between HFNs and LFNs (Confais et al., 2017). This suggests that the input-output gain of LFNs from DR afferents are under tonic suppression and/or that of HFNs are under tonic facilitation, both from external neural inputs (but (Matthews, 1999), see discussion). The fact that HFNs have regular firing rates (Fig. 6F) also supports this possibility because dominating excitatory input (weaker suppression or larger facilitation) results in regular spike intervals (Ponce-Alvarez, Kilavik, & Riehle, 2010; Stevens & Zador, 1998).

**Figure 7:**
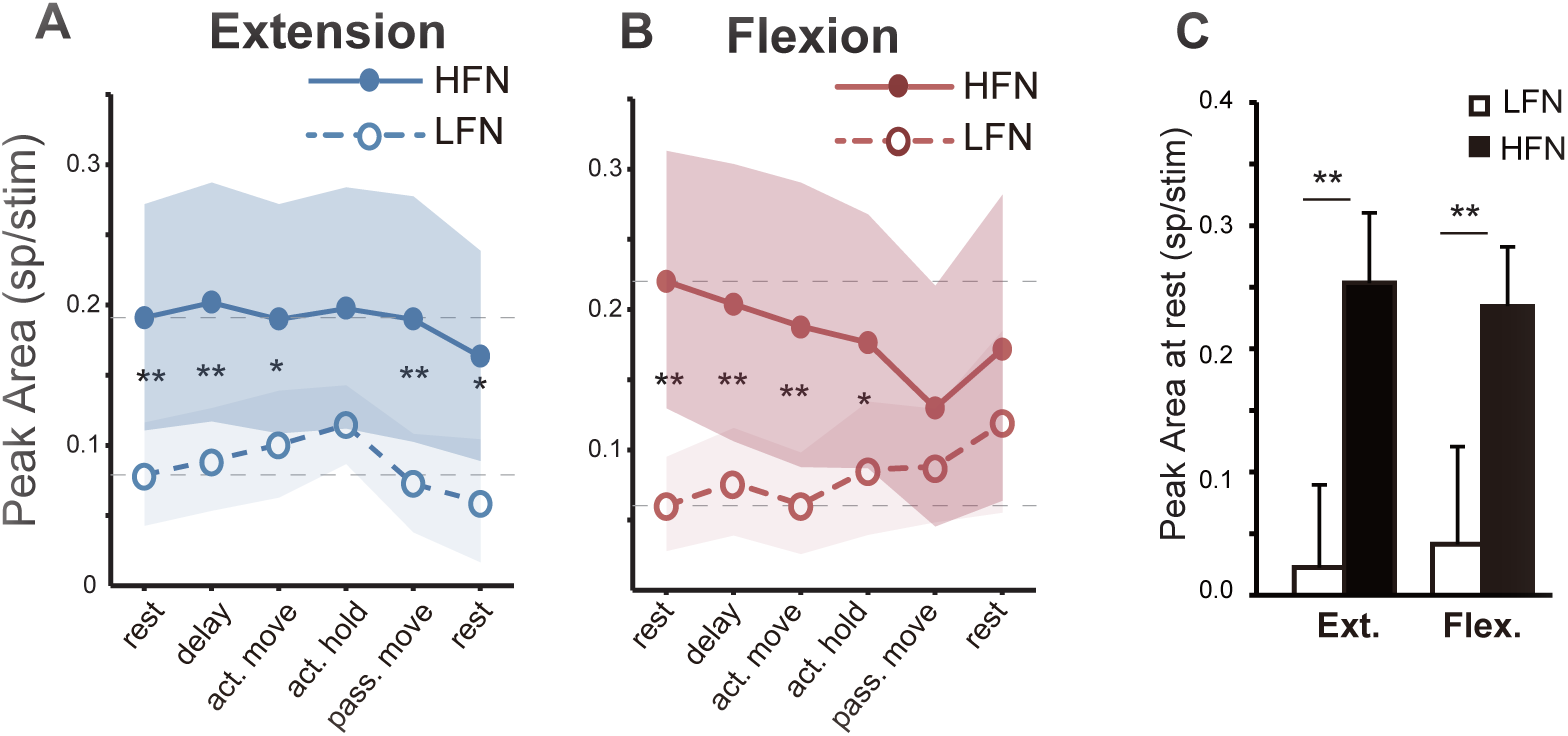
Population measure of evoked response in DR neurons. Median evoked peak area evoked in DR neurons during extension (A) and flexion (B) are presented for each behavioral epoch. Neurons showing larger firing rate during active flexion (HFN; filled) or active extension (LFN; open). Flexion trials are shown in red and extension trials are shown in blue. The shaded areas represent the interquartile range around each data point. Asterisks indicate that the median peak area in a specific epoch was significantly different between HFN and LFN groups (Wilcoxon rank sum test; *, *P* < 0.05; **, *P* < 0.01). (C) Mean peak areas during rest periods at the beginning of a task are compared between LFN (open) and HFN (filled) groups (two-sample t-test; *, *P* < 0.05; **, *P* < 0.01).

### SR neurons did not exhibit distinct subgroups

So far, we have found a distinct difference between two subgroups of DR neurons (HFNs and LFNs) in terms of their intrinsic properties, firing characteristics during wrist movements, and their underlying network properties (effectiveness of group I input). While it is likely that these findings are specific to DR neurons, it could also be interpreted as a shared, general property of spinal neurons receiving input from peripheral afferents. To examine this possibility, we repeated the analysis conducted in the DR neurons (Figs. 5–7) in the SR neurons (Fig. 8). First, by using the same criteria to dissociate HFNs and LFNs, we categorized SR neurons into two groups (F > E neuron vs. F < E neuron). Here, F > E neuron functionally corresponds to the HFNs, and F < E neurons to the LFNs of the DR neurons. As shown in Fig. 8, the findings described in Figs 5–7 could not be reproduced in the SR neurons. Both groups showed comparable firing patterns (except active extension, see (Seki et al., 2003, 2009)) (Fig. 8; A, B), spike width (C), base firing rate (D), firing irregularity (E), and responsiveness to SR afferent inputs (F, G, and H). These results suggest that the differences found between HFNs and LFNs were specific to spinal sensory neurons receiving proprioceptive afferent signals, and not to spinal sensory neurons in general.

**Figure 8:**
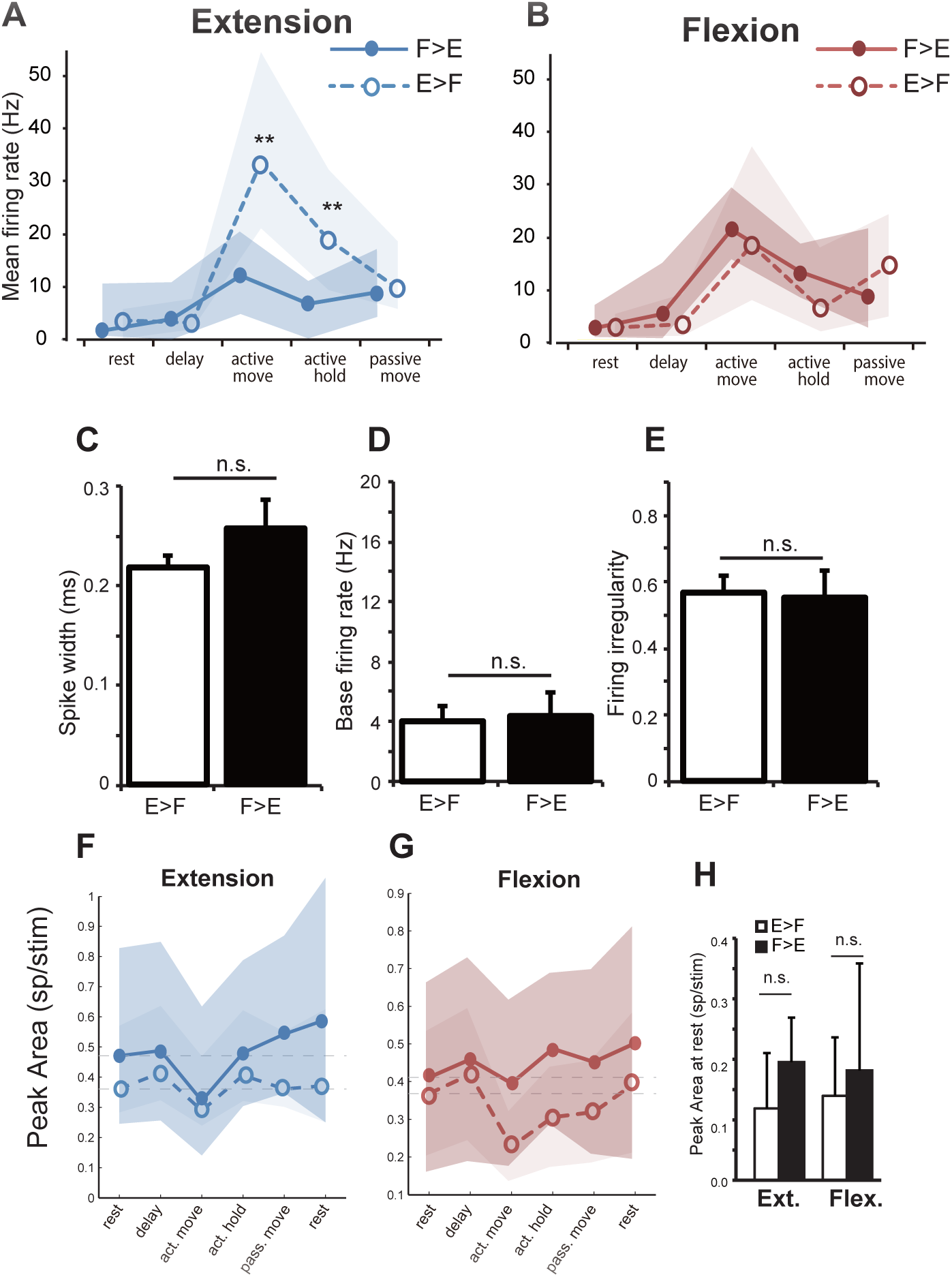
Summary of results in SR neurons. To compare the result of DR and SR neurons, the same analyses as in Figs. 5–7 were repeated in the SR neurons. For this purpose, SR neurons were subdivided in classes similar to those used for DR neurons: those with greater firing rates during flexion (F > E group, comparable with the HFNs in DR neurons) and those with greater firing rates during extension (F < E group, comparable with the LFNs in DR neurons). (A, B) population analysis of firing rate modulation (see Fig. 5). (C) spike-width, (D) base firing rate, (E) firing irregularity (see Fig. 6). (F, G, H) population measure of evoked response from SR nerve (see Fig. 7).

## Discussion

To elucidate the neuronal representation of proprioception during voluntary movements, we recorded the activity of spinal neurons receiving direct projections from spindle afferents in monkeys performing a simple wrist flexion-extension task. We found that proprioception may be represented in two functionally defined groups of spinal postsynaptic neurons in different ways: HFNs, with shorter spikes, slower background spike rates, and regular ISIs, seemed to be congruent with the activity of spindle-bearing muscles in terms of their firing modulation during the task. In contrast, LFNs, with longer spikes, high background activity, and irregular ISIs, seem to be incongruent with spindle activity. Therefore, we hypothesize that these two groups were sampled from distinct classes of spinal neurons with different intrinsic properties. In addition, HFN and LFN activity could be controlled by the different mode of synaptic inputs because the input-output gain from DR afferents to these neurons was different.

### Different intrinsic properties of HFNs and LFNs: their neuronal identity?

In the spinal cord, two types of neurons relay information from proprioceptors: ascending projection neurons and interneurons (Jankowska, 1992). In the cerebral cortex (Kaufman et al., 2010; Merchant et al., 2008; Miri et al., 2017; Mitchell et al., 2007; Mountcastle et al., 1969) or hippocampus (Henze et al., 2000), the duration of extracellular spike waveforms has been used to distinguish recordings from putative projection neurons (longer) vs interneurons (shorter). Although few comparable studies were found for the spinal cord, limited information suggests that ascending projection neurons could also be large (Todd, 2010; Todd, McGill, & Shehab, 2000). Consequently, we propose that LFN and HFN signals are predominantly recorded from ascending projection neurons and interneurons, respectively. Assuming that the majority of DR neurons are located in the anterior dorsal horn or the intermediate zone (laminae IV–VII) (Confais et al., 2017), we may further speculate about their potential identity.

Postsynaptic dorsal column neurons (PSDC neurons) are possible candidates for LFNs. First, they receive direct projections from Ia afferents (Jankowska et al., 1979; Maxwell, Koerber, & Bannatyne, 1985), mainly located around laminae IV to VI in monkeys (Rustioni, Hayes, & O’Neill, 1979). Second, they tend to include large cells (Enevoldson & Gordon, 1989) with fast conduction velocities (Jankowska et al., 1979), both suggesting spikes with longer durations. By contrast, the cell bodies of the major spinal ascending pathway are located too dorsally (spinotharamic) or ventrally (spinocervical), making them unlikely candidates for LFNs.

Several kinds of spinal interneurons of lamina IV-VII have been shown to be activated by muscle afferents with shorter latencies (Jankowska, 1992; Jankowska et al., 1979) and could be HFNs. They consist of excitatory and inhibitory neurons with different projection targets, including motoneurons, neurons of the contralateral side, or other interneurons. They vary in size and dendritic organization (Al-Mosawie, Wilson, & Brownstone, 2007). Therefore, even if HFNs more likely represent segmental interneurons, rather than ascending projection neurons, their identity is unclear.

### Congruent and incongruent representation of spindle signals in HFNs and LFNs

The firing rates of spindle afferents and that of their target neurons have been investigated in a few studies (Kolmodin, 1957; Osborn & Poppele, 1989); and it is known that dorsal spinocerebellar tract (DSCT) neurons encode information about muscle length fairly accurately (for review, see (Jankowska, 1992)). In the anaesthetized cat, Osborn and Poppele (Osborn & Poppele, 1989) recorded responses to muscle stretch, contraction, and the electrical stimulation of muscle-afferents in a population of DSCT neurons, that received direct projection from proprioceptors. They found that DSCT neurons were facilitated with remarkable similarity regardless of the type of stimulus used. Based on this, we could expect monotonic facilitation of DR neurons during lengthening of their spindle-bearing muscle (i.e. wrist flexion). In this sense, the congruent activity seen in HFNs (only 25% of DR neurons) could be expected, but the incongruent activity pattern found in LFNs (51% of DR neurons) was much less so. It is possible that the incongruent activity seen in LFNs is specific to the awake, behaving condition because neurons receiving projections from spindle afferent may also receive input from other sources (Baldissera, Hultborn, & Illert, 1981). These sources of convergent input may facilitate their target neurons in an agonistic or antagonistic way.

Fusimotor drive could be a potential source of the incongruent, comparatively larger, activity in LFNs during the shortening movement. Under alpha-gamma coupling, fusimotor drive to the spindle-bearing muscle of DR neurons should be larger during contraction, that is, extension, thus facilitating the activity of DR afferents. Were this effect dominant compared with the activity induced by simple mechanical stretching of the spindle-bearing muscles during movement, the observed incongruent activity could be expected. However, we think this possibility unlikely: First, spindle afferent recordings in cats (Loeb et al., 1985), and monkeys (Schieber & Thach, 1980), as well as microneurography studies in human subjects (al-Falahe et al., 1990a, 1990b), suggest that changes in muscle length can override the effect of alpha-gamma linkage. Second, it seems that fusimotor drive may not be selective enough to specifically influence a subset of spindles (e.g. those projecting axons to LFNs). Indeed, it has been shown that the pattern of fusimotor innervation is random and variable, both in terms of innervation point and action onto different types of intrafusal fibers (for review, see (Jankowska, 2015)). Therefore, we propose that Ia input to LFNs may be specifically modulated.

### Mechanisms underlying different input-output gains between HFNs and LFNs

We found that HFNs showed greater activity (Fig. 5) when their parent muscles were lengthened by wrist movements. Their high fidelity to the spindle input could be supported by higher input-output gain from Ia afferents. Indeed, the input-output gain from Ia afferent seems higher throughout movement (Fig. 7), both during flexion and extension. Consistent higher gain throughout the task could be advantageous for HFNs accurately encoding the length of spindle-bearing muscles induced by own movement or unpredictable perturbation. In contrast, the LFNs had much lower activity during flexion compared with extension (Fig. 5), even though input from spindle-bearing muscles increased. This suggests that LFNs could be uncoupled from their Ia input. This uncoupling between the afferent input to LFNs could be induced by a persistent lower input-output gain (Fig. 7). Under this condition, postsynaptic neurons are dissociated from their proprioceptive information encoded by Ia afferents. This might be advantageous in allowing LFNs to encode other inputs in a highly convergent system (Baldissera et al., 1981). For example, Ib, but not Ia, afferents in the DR nerve are highly active when spindle-bearing muscles are shortening (that is, during extension). By lowering the input-output gain of the Ia-LFNs synapse, LFNs could represent alternative inputs more faithfully.

We hypothesize that presynaptic inhibition could set a mode of coupling (HFN) or uncoupling (LFN) from Ia afferents. We previously showed that cutaneous input to spinal neurons could be modulated by presynaptic inhibition during voluntary movement (Seki et al., 2003, 2009). If the same mechanism also applies to the synapse from Ia afferents to spinal neurons, persistently lower presynaptic inhibition may cause coupling between Ia and HFNs, and higher presynaptic inhibition may cause uncoupling with LFNs. We speculate that convergent, antagonistic inputs to LFNs could be efficiently incorporated into postsynaptic systems according to their relevance to ongoing behavior. However, this implication should be made with considerable caution. Matthews (1999) proposed that input-output gains could be nonlinear when the excitatory drive to limb motoneurons is very low (Matthews, 1999). If this also applies to spinal interneurons, different input-output gains between HFNs and LFNs might be ascribed to differences in intrinsic neuronal properties. Therefore, confirmation of this hypothesis needs further studies evaluating the level of presynaptic inhibition of both HFNs and LFNs using more a direct method, such as excitability testing, (Ghez & Pisa, 1972; Seki et al., 2009; Tomatsu, Kim, Confais, & Seki, 2017).

### Proprioceptive representation in the spinal cord: interplay between cellular and network properties

As discussed, HFNs and LFNs probably belong to different classes of neurons (fast and slow firing cells), and could be differentially controlled by surrounding neuronal networks (low and high levels of presynaptic modulation). These differences in cellular and network properties may be functionally relevant, and not just a coincidence. HFNs presumably consist of local interneurons and their intrinsic properties (faster firing) could be advantageous in representing subtle changes in muscle length, even during rest. This faithful representation is crucial for some classes of segmental interneuron, for example reciprocal inhibitory interneurons, for proper balance between agonists and antagonists during behavior. The functions of these interneurons could be further secured by persistently lower input-output gains. Assuming that the majority of LFNs are ascending projection neurons, higher ascending systems may be unable to monitor the activity of Ia afferents through LFNs. This suggests that, in the ascending system, spindle activity is primarily represented by the direct pathway (Weinberg, Pierce, & Rustioni, 1990) and the secondary, indirect PSDC pathway may not be crucial for their proprioceptive representation, at least for the well-learned motor task in this study. Uncoupling between Ia afferents and LFNs could be beneficial for the ascending systems monitoring of information other than muscle length (e.g. muscle force signaled by the Gorgi tendon organ and Ib afferents).

Overall, proprioceptive inputs appear to be represented by two different neuronal classes of spinal neurons. Because these properties were not found in neurons receiving cutaneous afferents, this way of representing bodily information could be unique for proprioception. We suggest that while robust proprioceptive representation in the local interneurons are assured by activity congruent with that of spindle afferents, proprioceptive representation in ascending projection neurons could be more flexible and task dependent.

## Materials and methods

### Animals

Experiments were approved by the Institutional Animal Care and Use Committee of the National Institute for Physiological Sciences (NIPS) and the Institutional Animal Care and Use Committee of the University of Washington. We obtained data from two female *Macaca fuscata* (monkeys KO and OK), one male *Macaca fuscata* (monkey IS), and one male *Macaca nemestrina* (monkey KJ). Data from monkey KJ were recorded by one of the authors in the lab of Dr. Eberhard E. Fetz, and data from the other three animals were recorded at the NIPS in Okasaki, Japan. During training and recording sessions, each monkey sat upright in a primate chair (Nakazawa-Koumuten, Hongo, Tokyo, Japan) with his/her right arm restrained and elbow bent at 90°. The monkey’s hand was held in a cast, with fingers extended and the wrist in the mid-supination/pronation position. The cast holding the monkey’s hand was attached to a servomotor-driven manipulandum (Washington National Primate Research Center, Seattle, WA, USA) that measured the flexion–extension torque of the wrist and applied position-dependent (auxotonic) torque. The left arm was loosely restrained to the chair (Confais et al., 2017; Seki et al., 2003).

### Behavioral paradigm

The monkeys performed a wrist flexion–extension task with an instructed delay period (Confais et al., 2017; Prut & Fetz, 1999; Seki et al., 2003). The position of a cursor displayed on a video monitor in front of the monkey was controlled by wrist flexion– extension torque. Trials began with the monkey holding the cursor in a center target window corresponding to zero torque for 800–1500 ms (rest; Fig. 1A). Next, flexion and extension targets were shown to the left and right of the center target. One target was filled transiently for 200 – 500 ms (cue), indicating the movement to be performed at the end of the instructed delay period (delay), which was signaled by the disappearance of the center target (go). Trials were accepted only if no wrist movement occurred during the delay period (550 – 1750 ms).This delay period is a subject of redefinent relative to EMG onset, as described in below. Following a brief reaction time after the go signal, the monkey quickly (within 300 ms) moved (active move) the cursor to the desired target and held it in the target window for a period of 1000–1500 ms (active hold). The movements were performed against an elastic load applied by the servomotor. At the end of the active hold period the torque target disappeared and the center target reappeared (second go). After a second reaction time, the monkey relaxed her/his forearm muscles, allowing the servospring to return the wrist passively (passive move) to the zero-torque position (rest). After maintaining the cursor within the center target for 500–800 ms, the monkey was rewarded with apple sauce (reward) for successful trials. The spring constant of the servomotor was the same throughout this task sequence; thus, the auxotonic load applied to the monkey’s wrist remained constant. Therefore, any period when the wrist moved away from the zero-torque position was active movement (with auxotonic muscle contraction), and any period when the wrist moved toward the zero-torque position was passive movement (with little muscle contraction). A unique feature of this wrist flexion–extension task was that it allowed independent evaluation of a number of major components of voluntary movement in each behavioral epoch. During flexion trials, we evaluated the instructed delay period for flexion, active flexion movement, sustained flexion torque, and passive extension movement. During extension trials, we evaluated the instructed delay period for extension, active extension movement, sustained extension torque, and passive flexion movement. We investigated the activity of spinal neurons in these eight behavioral epochs.

### Surgical implants

After training, surgery was performed aseptically while the animals were under 1.5–3.0% sevoflurane anesthesia in 2:1 O_2_:N_2_O. Head stabilization lugs were cemented to the skull with dental acrylic and anchored to the bone *via* screws. A stainless-steel or plastic recording chamber was implanted over a hemi-laminectomy in the lower cervical vertebrae (C4–C7). Pairs of stainless-steel wires (AS631, Cooner Wire, Chatsworth, CA, USA) were implanted subcutaneously in 10–12 muscles (extensor carpi ulnaris, extensor carpi radialis, extensor digitorum communis, extensor digitorum-2,3, extensor digitorum-4,5 (ED45), flexor carpi radialis (FCR), flexor carpi ulnaris, flexor digitorum superficialis (FDS), palmaris longus, pronator teres, abductor pollicis longus, and brachioradialis) to measure the earliest onset of muscle activity. Three nerve cuff electrodes (Haugland, 1996) were implanted on the radial nerves for stimulation (Fig. 1B). One cuff was implanted in the muscle branch (DR nerve: approximately 2 cm proximal to the elbow joint). In addition, two cuff electrodes were separately implanted in the cutaneous branch (SR nerve) approximately midway between the elbow and the wrist (Seki & Fetz, 2003; Seki et al., 2009). Note that nerve cuffs were implanted in only the SR of monkey KJ. Nerve cuffs were also implanted in the median nerve in monkeys KO, IS, and OK, but the neural responses evoked by the stimulation to this nerve were not included in this paper because it contains indistinguishable cutaneous and muscle afferents (see (Confais et al., 2017)).

### Classification of spinal neurons

Glass-insulated tungsten (Alpha Omega, Nazareth, Israel) or Elgiloy microelectrodes (impedance = 0.8 to 1.4 MΩ) were used for recording neural activity. At the beginning of each electrode penetration, cord surface potentials were monitored and threshold currents that evoked an incoming volley from the SR and DR were measured on each recording day. Subsequently, spinal neuronal single-unit responses were examined by biphasic constant-current pulse (100 μs/phase) stimulations at a constant frequency of 1–3 Hz.

For each neuron and stimulated nerve, we generated a PSTH, aligned to the timing of the stimulation pulses (Figure 1D, bottom). Each action potential was isolated based on its waveform (MSD, Alpha Omega), and PSTHs were obtained for each neuron (Fig. 1D). The PSTH included the data from 50 ms before to 30 ms after the stimulation pulse, with a bin size of 0.1 ms. The method we used to compute the peak amplitude was similar to that employed by Seki et al. (2003), as follows: The baseline firing rate was first computed as the mean bin height in the 50 ms epoch preceding the stimulation pulse. The peak onset was defined as the time at which the PSTH following the stimulation pulse crossed two standard deviations above the average of the baseline firing rate. Similarly, the peak offset was defined as the time at which the PSTH achieved the same criterion a second time. To ensure the algorithm did not detect a local maximum instead of a more visible peak, we used a set of additional ad hoc criteria: First, the peak onset had to occur inside a specific time window (from 0–5 ms after DR or M stimulation, and from 3–13 ms following SR stimulation). Second, the peak had to be > 70% of the highest bin from the beginning of the PSTH until the end of the detection window. If the detected peak failed to satisfy these ad hoc criteria, the next detected peak was selected, and so on. The peak area was computed as the sum of the bins above baseline firing rate during the peak duration (i.e., the grey area in Figures 1C and 4A-C). Thus, the peak area can be understood as the mean number of spikes above baseline evoked by each stimulation pulse during the peak time window. Stimulus current was set at 1–1.2 times the threshold for DR to activate primarily the Ia afferents, and the current was twice the threshold for SR. The segmental response latency was calculated from the entry of the volley into the spinal cord (the first positive peak of incoming volleys recorded in the cord surface potential (Eccles, Fatt, & Landgren, 1956)) to the onset of the PSTH peak. A central latency of less than 1.5 ms was adopted as a criterion for monosynaptic linkage from the SR or DR, and thus used to classify the neurons with short latency input (putative first-order interneurons) from each nerve (Egger, Freeman, Jacquin, Proshansky, & Semba, 1986; Moschovakis, Sholomenko, & Burke, 1991; Seki & Fetz, 2003). If neurons showed a large “motor unit” signature in the spike-triggered average of the unrectified EMG with only 50 spikes (Maier, Perlmutter, & Fetz, 1998) and at least one of 10–12 muscles recorded, they were classified as putative motoneurons and were excluded from the data set. Therefore, our data set consisted of putative first-order interneurons from given primary afferent (e.g., segmental interneurons, ascending projection neurons, or propriospinal neurons) but contains no putative motoneurons innervating the wrist and finger muscles.

### Characterization of interneurons based on their activity during wrist movement

After classifying putative first-order interneurons based on the segmental latency from the SR and DR, the epoch-dependent modulation of firing rates during the wrist flexion–extension task was evaluated. For this purpose, each behavioral epoch was redefined in terms of wrist torque during the task (F–E torque) as follows: Movement onset (both volitional and passive) was defined as an instance of rapid and steady change in the filtered derivative of F–E torque (50 Hz low-pass); and movement offset was defined as an instance of the first local peak or the first point of zero-crossing in the filtered derivative of F–E torque (5 Hz low-pass) after an instance of movement at top velocity.

After defining the movement onset and offset, we defined five movement-related epochs: 1) rest, 500-ms interval before onset of cue signal; 2) delay, time from cue onset until either onset of the go signal or 300 ms before movement onset, whichever occurred first (Prut & Fetz, 1999); 3) active move, time from onset of volitional movement to offset of the movement; 4) active hold, time from offset of volitional movement to onset of return signal; and 5) passive move, time from onset of passive return movement to offset of the movement.

Next, the mean firing rate was calculated for each behavioral epoch of each neuron. To eliminate the effect of stimulus-induced events (both responses and stimulus artifacts), the period 5 ms before to 90 ms after each stimulus was excluded from this calculation.

### Compiling DR and SR evoked responses for each behavioral epoch

In addition to the mean firing rate, we measured the evoked response of each neuron during each behavioral epoch (see also (Confais et al., 2017)). For each DR and SR neuron, the mean evoked response was computed by pooling all the stimulation pulses recorded during the task to maximize the signal-to-noise ratio. We then used the onset and offset points of the mean evoked response to compute the peak area evoked in each task epoch (that is, using the stimulation pulses applied during a specific epoch). We restricted the analysis to those behavioral epochs in which a sufficient number of stimulus pulses were applied, to obtain an unbiased estimate of the evoked response area. The minimum number of stimulation pulses was thus arbitrarily set at 16 stimulations per epoch, as a compromise between the reliability of the peak area and the size of the database. Therefore, the number of analyzed neurons differs between epochs.

### Intrinsic firing properties of DR and SR neurons

To evaluate the intrinsic properties of recorded neurons, we measured 1) spike duration, 2) firing regularity, 3) resting spontaneous firing rate, and 4) recording depth.

We calculated the duration of spontaneous spikes from the negative trough to the next positive peak (Henze et al., 2000; Kaufman et al., 2010; Mitchell et al., 2007; Vigneswaran, Kraskov, & Lemon, 2011). This measure was taken from the averaged spike waveform (Fig. 6A inset). We averaged a mean number of 3887 ± 1058 spikes. To evaluate firing regularity/irregularity, we calculated the Lv of ISIs (Abbasi et al., 2017; Mochizuki et al., 2016; Shinomoto et al., 2003). The Lv measure was chosen because it represents local spike train variability between two successive ISIs, thus the measure is theoretically independent from firing rate (e.g. trial-to-trial variability). Lv was given as (Shinomoto et al., 2003),

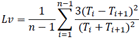

where T*i* is the duration of the *i*th ISI and *n* is the number of ISIs. Lv was calculated using the ISIs occurring in all behavioral epochs (Fig. 1A). ISIs around the stimuli to either DR or SR were eliminated from this calculation.

To evaluate resting spontaneous firing rate, we used the mean firing rate calculated during the rest period for both flexion and extension. Finally, the recording depth of each neuron within the spinal cord was evaluated by measuring the distance between each recording site and the location of the first neuron that we encountered in each penetration (these were expected to be located close to the posterior edge of the dorsal horn) (Confais et al., 2017; Takei & Seki, 2010).

### Statistics

We used Wilcoxon signed rank tests and Student’s t-tests to compare differences in means. R (R Foundation for Statistical Computing, Vienna, Australia) was used for the statistical analyses. We used the chi-square test to compare ratios between nerves or neuron groups. A *p-*value < 0.05 was considered to indicate statistical significance.

## Acknowledgements

We thank Nobuaki Takahashi (National Institute for Physiological Sciences) for technical assistance, Eberhard E. Fetz and Steve I. Perlmutter (University of Washington, Seattle, USA) for their permission to use the data on spinal interneurons with superficial radial input. This work was supported by National Institutes of Health grants NS 12542, NS36781 and RR00166, and Human Frontiers Science Program grant LT0070/1999-B, Grants-in-Aid 18020030, 18047027, and 26120003 for Scientific Research on Priority Areas: “Mobilligence,” “System Study on Higher-order Brain Function,” and Innovative Areas “Understanding Brain Plasticity on Body Representations to Promote their Adaptive Functions,” respectively, from the Ministry of Education, Culture, Sports, Science and Technology of Japan (MEXT) and the Japan Science and Technology Agency Precursory Research for Embryonic Science and Technology programs. The current address of TT is Centre for Neuroscience Studies, Queen’s University, Kingston, Ontario, Canada K7L 3N6.

## Competing Interests

The authors declare no competing financial interests.

